# Radial askew endodermal cell divisions reveal IRK functions in division orientation

**DOI:** 10.1101/2023.03.31.534810

**Authors:** R. M. Imtiaz Karim Rony, Roya Campos, Patricio Perez-Henriquez, Jaimie M. Van Norman

## Abstract

Oriented cell divisions establish plant tissue and organ patterning and produce different cell types; this is particularly true of the highly organized Arabidopsis root meristem. Mutant alleles of *INFLORESCENCE AND ROOT APICES RECEPTOR KINASE* (*IRK*) exhibit excess cell divisions in the root endodermis. IRK is a transmembrane receptor kinase that localizes to the outer polar domain of these cells, which suggests directional signal perception is necessary to repress endodermal cell division. Here, a detailed examination revealed many of the excess endodermal divisions in *irk* have division planes that specifically skew towards the outer lateral side, therefore we termed them ‘radial askew’ divisions. Expression of an IRK truncation, lacking the kinase domain, retains polar localization and rescues these radial askew divisions, but the roots exhibit excess periclinal endodermal divisions. Using markers of cell identity, we show that the daughters of radial askew divisions transition from endodermal to cortex identity similar to those of periclinal divisions. These results extend the requirement for IRK beyond repression of cell division activity to include cell division plane positioning. Based on its polarity, we propose that IRK at the outer lateral endodermal cell face participates in division plane positioning to ensure normal root ground tissue patterning.

## INTRODUCTION

In multicellular eukaryotes, cell division produces diverse cell types and increases cell number for body plan elaboration and growth. Cell divisions are typically categorized as proliferative (symmetric) and formative (asymmetric) . Proliferative cell divisions are typically physically symmetrical, producing daughter cells of the same size and cell fate. In contrast, formative cell divisions can be physically symmetrical or asymmetrical but the daughter cells acquire distinct identities. Due to the cell wall, the relative position of individual plant cells is fixed in space, as such, previous division orientations are ‘recorded’ by the placement of the walls (Facette et al., 2018; Rasmussen and Bellinger, 2018). The orientation of plant cell divisions is informed by extracellular cues and intracellular polarized proteins and is crucial for plant cell fate determination and tissue organization (Rasmussen et al., 2011; Van Norman, 2016; Wallner, 2020; Hartman and Muroyama, 2023). Because the developmental trajectory of a daughter cell is influenced by positional information (van den Berg et al., 1995; Marhava et al., 2019), division plane orientation is a key developmental decision.

The orientation of plant cell divisions is defined by how the new cell wall is positioned relative to the surface of the organ (Livanos and Müller, 2019). Accordingly, periclinal cell divisions are oriented parallel to the surface and anticlinal cell divisions are oriented perpendicular to the surface. For anticlinal cell divisions, the division plane can be aligned with the organ’s longitudinal or transverse axis (Figure 1A, B); therefore, we distinguish between longitudinal anticlinal and transverse anticlinal cell divisions. In the root, periclinal cell divisions are typically formative as they generate additional layers of distinct cell types (De Smet and Beeckman, 2011; Pillitteri et al., 2016). Whereas, anticlinal cell divisions are often proliferative, generating more cells of the same type. Normal tissue and organ patterning require the specific orientation of proliferative and formative cell divisions relative to the plant body axes. Yet our understanding of how division plane orientation is precisely controlled and how it is coupled with cell division frequency remains limited.

The highly stereotypical cellular organization of the *Arabidopsis thaliana* root makes it an ideal system to investigate how the frequency and orientation of cell division contribute to tissue/organ patterning. The root ground tissue (GT), is a good example of how a specific series of oriented formative cell divisions give rise to distinct cell types, the cortex and endodermis. First, the GT stem cell, the cortex/endodermal initial (CEI) (Figure 1A, B), undergoes a formative, transverse anticlinal division to produce the CEI daughter cell (CEID). This daughter cell then undergoes a formative, periclinal division to produce endodermis towards the inside and cortex cells towards the periphery of the root (Figure 1B) (Dolan et al., 1993; Scheres and Benfey, 1999)). As plants mature, endodermal cells can undergo another periclinal division to produce another cortex layer called middle or secondary cortex (Paquette and Benfey, 2005; Cui, 2016). After their formation, cortex and endodermal cells proliferate through symmetrical, transverse anticlinal divisions, producing more cells in their respective longitudinal files. Symmetrical, longitudinal anticlinal divisions (LADs) can also occur producing more GT cells in the root’s radial axis (Figure 1A), however, these divisions are rarely observed in young wild type roots and, as a result, eight of each GT cell type, including the CEI, are most often observed around the stele (Dolan et al., 1993; Scheres and Benfey, 1999). Deviations in cell division planes lead to irregular daughter cell shapes allowing the consequences of misoriented root cell divisions to be followed in time and space.

Strict regulation of GT cell division in the root’s radial axis is disrupted in mutant alleles of *IRK* (*INFLORESCENCE AND ROOT APICES RECEPTOR KINASE*), which exhibit excess endodermal longitudinal anticlinal and periclinal cell divisions (Campos et al., 2020). These excess cell divisions coincide with promoter activity of *CYCLIN D6;1* (Campos et al., 2020), a specific D-type cyclin, which is associated with formative, but not proliferative, GT cell divisions (Sozzani et al., 2010). Yet, no increase in proliferative, transverse anticlinal divisions was observed in *irk-4*, suggesting IRK specifically represses cell divisions that produce more cells in the root’s radial axis (Campos et al., 2020). We propose that IRK specifically represses an endodermal cell division program that widens the GT and that longitudinal anticlinal and periclinal endodermal cell divisions are developmentally regulated downstream of IRK.

IRK encodes a transmembrane receptor kinase of the Arabidopsis RECEPTOR-LIKE KINASE (RLK) super family (Shiu and Bleecker, 2001; Shiu and Bleecker, 2003) and accumulates to the outer polar domain of the endodermal plasma membrane (PM). Although *IRK* is expressed in several root cell types, endodermal-specific expression of *IRK* is sufficient to rescue the cell division defects (Campos et al., 2020; Rodriguez-Furlan et al., 2022). Unexpectedly, IRK maintains its polar localization when the intracellular kinase domain is missing and this IRK truncation remains largely functional, rescuing the endodermal LADs but not the periclinal divisions in *irk-4* (Rodriguez-Furlan et al., 2022). Polar accumulation of IRK to the outer lateral endodermal cell face suggests it perceives an extracellular cue that originates peripheral to the endodermis and is required to maintain GT organization and cell number in the root’s radial axis. However, the molecular details of IRK function remain unclear.

Here, we closely examined the excess endodermal divisions in *irk* mutants and identified a distinct cell division orientation defect. Unexpectedly in *irk-4*, a null allele, the majority of endodermal cell divisions previously characterized as periclinally oriented, instead, have outwardly skewed division planes. We defined these as ‘radial askew’ endodermal divisions, because these oblique divisions consistently produce small, abnormally shaped cells peripheral to the endodermis. In the weaker *irk-1* allele, roots have fewer radial askew endodermal cell divisions, but have excess longitudinal anticlinal and periclinal divisions. Therefore, while both *irk* alleles have excess GT cells in the radial axis, the orientation of the excess cell divisions is different. Additionally, the IRK intracellular truncation rescues the radially askew endodermal divisions in *irk-4*, but excess periclinal divisions are present, phenocopying *irk-1*. These results expand the function of IRK beyond repression of endodermal cell division activity showing it is also required for cell division plane orientation. Given its polar accumulation in the PM, IRK may participate in division plane orientation through physical involvement with cell division machinery or through perception of directional, positional information. We propose a model whereby the presence of IRK at the outer polar domain occludes that endodermal cell face for division plane selection or cell plate attachment.

## RESULTS

### Many endodermal cell divisions in *irk-4* are abnormally oriented

Close examination of endodermal cell division orientation in *irk-4* revealed that divisions we had classified as periclinal were not actually oriented parallel to the root’s surface. If a periclinal division is considered in 2-dimensions (2D) in the longitudinal axis, the new cell plate attaches to sites to opposite root/shootward endodermal faces within a cell file. However, in *irk-4*, we observed abnormal divisions where cell plates attach to adjacent endodermal faces - either the shoot- or rootward face and the outer lateral faces (Figure 1C-E). These abnormal, oblique division planes resulted in new cell walls that often appeared periclinally oriented when viewed only in longitudinal optical sections. However, in 3D, as observed in a series of longitudinal and transverse optical sections (in confocal z-stacks (Campos and Van Norman, 2022)), key differences between true periclinal and these abnormal divisions become apparent. Periclinal endodermal divisions produce daughter cells of nearly equal size, whereas the abnormal divisions in *irk-4* result in unequal sized daughter cells with the smaller, prism-shaped cell peripherally positioned. Intriguingly, we did not observe any abnormal divisions oriented toward the inner lateral side of the endodermis, indicating division planes in *irk-4* are not randomly misoriented. Given the specific orientation of these divisions, we termed them radial askew endodermal divisions.

To assess the frequency of radial askew endodermal divisions, we examined a set of confocal images (Rodriguez-Furlan et al., 2022) to specifically parse radial askew from periclinal endodermal divisions. This revealed that 88.8% of the endodermal divisions in *irk-4* classified as periclinal (n = 224 divisions) were actually radial askew divisions. Thus, there was no significant difference in the number of periclinal divisions in *irk-4* compared to wild type (Wt) (Figure 1F). In young Wt roots, periclinal divisions are somewhat rare and 5.1% of the divisions classified as periclinal (n = 59 divisions) were found to be radial askew. In *irk-1*, a partial loss of function allele, we found that 15.6% of divisions classified as periclinal (n = 154 divisions) are radial askew oriented. Despite this, *irk-1* roots have significantly more periclinal divisions than Wt (Figure 1F). These analyses revealed key phenotypic differences between the *irk* alleles: *irk-4* has substantially more radial askew and longitudinal anticlinal divisions than Wt or *irk-1*, whereas *irk-1* has significantly more periclinal and longitudinal anticlinal divisions that Wt. Because *irk-1* has many fewer radial askew endodermal divisions than *irk-4*, these divisions appear to be a specific phenotype observed upon complete loss of IRK function. These observations indicate that IRK operates not only to repress endodermal cell division activity but also functions in division orientation.

### The IRK kinase domain is dispensable for its role in cell division orientation

We previously showed that GFP fused to a truncated version of IRK missing its kinase domain (IRKΔK-GFP) was polarly localized in the endodermis (Figure 1G) and, while it fully rescued the *irk-4* endodermal longitudinal anticlinal division (LAD) phenotype, it did not rescue the periclinal division phenotype (Rodriguez-Furlan et al., 2022). This is reminiscent of the *irk-1* root phenotype. As *irk-1* contains a T-DNA insertion near the beginning of the region coding for the kinase domain (Campos et al., 2020), we predict that production of a partially functional truncation results in the milder *irk-1* cell division phenotype. With the detection of novel phenotypic consequences of *irk* loss of function and differences between the *irk* alleles, we investigated whether IRKΔK-GFP could rescue the radial askew cell division phenotype of *irk-4*.

We found that *irk-4* roots expressing the full-length IRK-GFP or IRKΔK-GFP showed rescue of endodermal LAD and radial askew phenotypes (Figure 1G, H). However, the *irk-4* IRKΔK-GFP roots had significantly more periclinal endodermal divisions than Wt or *irk-4* IRK-GFP plants, making them phenotypically similar to *irk-1*. This indicates presence of IRKΔK at the outer polar domain is sufficient to repress longitudinal anticlinal and radial askew endodermal divisions in *irk-4*, but leads to excess periclinal divisions. Thus, although excess endodermal divisions are present in *irk-4* IRKΔK-GFP roots, they are predominantly periclinally oriented, which is more typical for endodermal formative divisions (Paquette and Benfey, 2005; Cui, 2016). These observations suggest the IRK kinase domain is largely dispensable in endodermal division plane orientation, but is involved in repression of endodermal cell division activity.

### Radial askew endodermal cell divisions in *irk-4* coincide with *CYCD6;1* promoter activity

In Wt roots, formative GT cell divisions are associated with promoter activity of a specific D-type cyclin, *pCYCD6;1*, this includes divisions of the initial cells and periclinal endodermal divisions, which generate middle cortex (Sozzani et al., 2010). In *irk* mutants, excess endodermal LADs also coincide with *pCYCD6;1* activity (Campos et al., 2020). This putatively links longitudinal anticlinal and periclinal endodermal divisions through some shared regulatory mechanism(s) downstream of IRK function and suggests IRK specifically represses an endodermal cell division program that operates in the root’s radial axis.

With the addition of radial askew divisions to the population of excess endodermal divisions in *irk-4*, we conducted a detailed examination of *pCYCD6;1:erGFP* expression in *irk* and Wt roots grown on 0.2x MS, which aggravates *irk* root phenotypes (Campos and Van Norman, 2022; Goff et al., 2023). Similar to our previous report, *pCYCD6;1* activity is substantially increased in the *irk* endodermis compared to Wt. We quantified abnormal endodermal cell divisions together with expression of *pCYCD6;1:erGFP* and mapped this information in the root’s longitudinal axis (Figures 2, 3, and S1). In Wt roots, endodermal LADs occur much less frequently than in *irk* and the daughter cells are rarely associated with *pCYCD6;1* activity (Figure 2A-D). Under these conditions, periclinal divisions occur in Wt at this age (6 days post-stratification) and, as expected, often associate with *pCYCD6;1* activity (Figure S1A-D). In *irk-1*, there are more endodermal LADs than in Wt with daughter cells frequently showing *pCYCD6;1* activity. *irk-4* roots have endodermal LADs at nearly every position in the longitudinal axis and, with rare exception, these daughter cells show *pCYCD6;1* activity (Figure 2C, D). Like in Wt, periclinal endodermal divisions in *irk* alleles most often associate with *pCYCD6;1* activity (Figure S1C, D). These results reveal extensive excess endodermal LADs in *irk* mutants and confirm their strong association with *pCYCD6;1* activity.

Radial askew endodermal cell divisions are also associated with *pCYCD6;1* activity (Figure 3). In the rare instances when radial askew divisions are observed in the Wt endodermis, they are associated with *pCYCD6;1* activity (Figure 3A-C). This association was also observed in *irk* mutants, where there are numerous radial askew endodermal divisions (Figure 3B, C). In Wt and *irk-1*, radial askew endodermal divisions begin to occur five or more cells above the quiescent center (QC), however in *irk-4*, radial askew divisions occur much more frequently and closer to the QC (Figure 3C). Like endodermal LADs in *irk*, the daughter cells of radial askew divisions nearly always show *pCYCD6;1* activity. Altogether, our endodermal cell division maps reveal extensive excess endodermal cell divisions in the *irk* mutants, particularly *irk-4*, where the daughter cells of LADs and radial askew divisions consistently show *pCYCD6;1* activity. The association of *pCYCD6;1* activity with each of the excess endodermal cell division orientations in *irk* further demonstrates the requirement for IRK in repression of cell division activity in the endodermis.

### Peripheral daughter cells of radial askew endodermal divisions often express a marker of cortex identity

Endodermal periclinal, longitudinal anticlinal, and radial askew cell divisions all result in an increased number of GT cells in the radial axis of *irk* roots. LADs produce more endodermal cells around the stele and periclinal divisions produce a third GT layer, the middle cortex. We expect that the daughters of endodermal LADs will maintain endodermal identity, while those of periclinal divisions will transition to cortex identity (Figures 4, S2, and S3). As endodermal radial askew cell divisions produce peripherally located, prism-shaped cells of unknown fate, we examined reporter gene expression for endodermal and cortex cell identity (Figure 4). We first examined *SCARECROW* promoter activity (*pSCR:erGFP*), which is expressed in endodermal cells throughout the root (Di Laurenzio et al., 1996; Wysocka-Diller et al., 2000). In Wt, 43% (3 of 7) of peripheral daughters of radial askew divisions show *pSCR* activity. In *irk-1* and *irk-4*, 94% (25 of 26) and 60% (176 of 293) of these daughter cells show *pSCR* activity (Figure 4A-C), respectively. Thus, regardless of genotype, not all peripheral daughters of radial askew endodermal divisions show *pSCR* activity, implying they do not maintain endodermal identity.

Next, we examined activity of the *CORTEX2* promoter driving nuclear localized-yellow fluorescent protein (*pCO2:nlsYFP*), which is expressed in cortex cells in the root meristem (Heidstra et al., 2004; Paquette and Benfey, 2005). In Wt roots, cells immediately peripheral to the endodermis have cortex identity (Figure 1B), whether those cells are derived from periclinal CEID or endodermal divisions. Radial askew divisions were very infrequent in Wt and *irk-1* expressing *pCO2:nlsYFP*. The single peripheral radial askew daughter in Wt did not express *pCO2* and only one of four did in *irk-1*. In *irk-4*, ∼21% (46 of 216) of the peripheral daughters of radial askew divisions exhibited *pCO2* activity (Figure 4D-F). Comparison of the activity of *pSCR* and *pCO2*, particularly in *irk-4*, suggests that peripheral daughters of endodermal radial askew divisions gradually acquire cortex cell fate similar to what is observed for the peripheral daughters of endodermal periclinal divisions.

To more directly investigate this, we examined *irk-4* roots simultaneously expressing *pSCR* and *pCO2* reporters. Using linear unmixing, the daughters of endodermal longitudinal anticlinal, periclinal, and radial askew divisions were examined (Figure 5A-D). The daughter cells of endodermal LADs exclusively showed *pSCR* expression as expected, whereas the daughter cells of periclinal and radial askew divisions showed expression of either or both of these markers (Figure 5B). This suggests the daughter cells of radial askew and periclinal divisions progress along a similar developmental trajectory and supports the idea that radial askew divisions are formative divisions. Moreover, our observations suggest that *pCO2* activity is progressively turned on in more distal peripheral daughters in files of these prism-shaped cells (Figure 5C). Lastly, peripheral radial askew daughter cells more frequently show simultaneous *pSCR* and *pCO2* activity, suggesting acquisition of cortex cell identity progresses more slowly, than in the peripheral daughters of periclinal divisions (Figure 5D). Overall, we find that peripheral radial askew daughters acquire cortex identity.

### Radial askew and periclinal endodermal divisions often co-occur in the root’s longitudinal axis

Typically root cell divisions bisect cells strictly parallel or perpendicular to the root surface. When considered in 2D in the transverse axis, periclinal endodermal divisions occur when a new cell plate attaches to the circumferential endodermal faces and LADs occur when the new cell plate attaches to the outer and inner endodermal faces. In each division, the pair of resulting daughter cells of each division are similarly sized, but periclinal divisions are formative and produce a new cell layer with a distinct cell fate (middle cortex, Figure 1A-C). In contrast, the daughters of endodermal LADs remain endodermal. Radial askew endodermal divisions are distinct, with a division plane oblique to the root surface that creates daughter cells of different sizes, with the smaller prism-shaped cell consistently positioned peripherally. When considered in 2D in the transverse axis, these divisions occur when a new cell plate attaches to the outer lateral endodermal face and an adjacent circumferential endodermal face. Given the positions of cell plate attachment (Figure 1C), radial askew divisions could simply be misoriented longitudinal anticlinal or periclinal endodermal cell divisions. To address this we determined whether endodermal cell division orientations could be spatially correlated.

Within individual cell files, we located endodermal cells that had undergone radial askew division and assessed the number of instances in which the adjacent shootward and/or rootward cell(s) had undergone a periclinal or longitudinal anticlinal division or had not divided at all (immediately adjacent radial askew divisions were not counted, Figure 5E). In Wt, the rare radial askew endodermal divisions (2 instances) had shootward and/or rootward neighbors that had divided periclinally or not at all (Figure 5F). In *irk-1*, radial askew endodermal divisions were adjacent to a periclinal division >60% of the time. Otherwise, the neighbor had not divided at all. In *irk-4*, radial askew divisions were most often adjacent to endodermal cells that had not divided and to periclinally divided cells just over 30% of the time. Unexpectedly, radial askew divisions in *irk-4* were very rarely flanked by LADs (Figure 5F). Given that endodermal radial askew divisions are very rare in Wt, their co-occurrence with periclinal division is notable. Likewise in *irk-4*, it is striking that radial askew divisions are usually adjacent to endodermal cells that have not divided or have undergone a periclinal division, given that endodermal LADs in *irk-4* are extremely numerous. These results suggest that, regardless of genotype, endodermal radial askew divisions are most often adjacent to cells that have divided periclinally or not divided at all. Given that endodermal radial askew daughter cells are peripherally located, transition to cortex identity, and their neighbors are more likely to have undergone a periclinal division, we propose radial askew divisions are abnormal GT formative divisions.

## DISCUSSION

We have identified excess and obliquely skewed endodermal cell divisions in *irk* mutants, a novel aspect of the mutant phenotype that reveals IRK functions to repress endodermal cell division activity and is required for cell division orientation. Typically, endodermal cell divisions bisect the cell into similarly sized daughters, however, the divisions we define as radial askew have oblique division planes creating small prism-shaped cells. Of particular surprise is the invariable positioning of these smaller daughter cells towards the outer lateral side -nearer the cortex. If there were random misorientation of endodermal divisions in *irk*, we would expect that approximately half would be oriented towards the inner lateral side, such that the smaller daughter cells formed nearer the pericycle. Because this was not observed in any of our experiments, there is no support for random misorientation of endodermal cell division in *irk*; instead, these divisions have a specific orientation. This is consistent with a hypothesis where positional information orients cell division planes away from the lateral endodermal cell faces in Wt, but in *irk* mutants, information is missing and division planes frequently contact the outer lateral face.

The specific orientations of the excess divisions among *irk* mutants suggest endodermal division activity and orientation are linked and tightly controlled. The role of IRK in this control is particularly compelling when the spatial relationship between the *irk* cell division orientation defects and IRK polar accumulation are considered (Figure 6A, B). In the endodermis of *irk-1* or *irk-4 IRKΔK-GFP* roots a portion of IRK is present at the outer lateral domain of the PM and there are fewer radial askew and LADs, but more periclinal divisions than in *irk-4*. In those genotypes, excess endodermal cell divisions are present but the division planes contact the outer lateral endodermal face infrequently. Whereas in *irk-4*, the putative null allele, excess endodermal divisions are very frequently oriented such that the division plane contacts the outer lateral face (Figure 6C). Therefore, the presence of IRK or IRKΔK at the outer lateral endodermal face has a negative, potentially repellent, influence on division plane positioning.

To instruct cell division plane orientation, a polar-localized protein would need to influence positioning of the cell division machinery. In plants, this includes the mitotic spindle and two plant-specific cytoskeletal structures, the preprophase band, which marks the future division site, and the phragmoplast, which assembles and guides cell plate formation (Rasmussen et al., 2013; Smertenko et al., 2017). Many studies have investigated the mechanisms underlying how a new cell plate is attracted to the marked division site. However, our findings are consistent with a mechanism that repels cell plates from particular sites. We predict dueling attractive and repulsive cues would increase the precision and robustness of the cell division orientation process. For instance, in the *ton recruiting motif* (*trm) 6*,*7*,*8* triple mutant the preprophase band is not detectable, yet the defects in root cell division orientation and stomata patterning are modest (Schaefer et al., 2017). This hints that other mechanisms exist to position division planes and there is increasing evidence that polarized proteins can directly and/or indirectly position division planes via linkage with the cytoskeleton. For example, during stomatal development, BASL/BRX family polarity domain depletes cortical microtubules at particular positions ensuring the preprophase band forms outside the site marked by this polarity domain (Muroyama et al., 2020; Muroyama et al., 2023). Identification of the molecular links between IRK and the mechanics of cell division orientation and/or cytokinesis are key avenues of future investigation.

We found radial askew endodermal divisions often spatially correlate with periclinal divisions and their peripheral daughters progressively acquire cortex identity, suggesting that they are formative in nature. Formative plant cell divisions often must traverse a longer path than proliferative divisions, which typically occur along the shortest path that divides the cell equally (Besson and Dumais, 2011; Facette et al., 2018; Rasmussen and Bellinger, 2018); indeed, the path of an endodermal periclinal division is the longest in this cell type. Therefore, a simple conclusion is that radial askew endodermal divisions are just misoriented periclinal divisions, this would suggest IRK is needed to maintain division orientation parallel to the outer lateral face during ‘long path’ endodermal formative divisions. However, contemplating IRK function only in the context of maintaining the orientation of formative, periclinal endodermal divisions due to the long division path fails to account for the excess endodermal LADs, which can also be considered ‘long path’ divisions and are a key attribute of the *irk* phenotype.

Moreover, radial askew divisions are often flanked by endodermal cells that have not divided at all. Comparing the instances of radial askew endodermal divisions in *irk-4* flanked by no division, an LAD, or a periclinal division to the average number of radial askew divisions per root, reveals that radial askew divisions occur in consecutive stretches along endodermal cell files. If they were simply misoriented periclinal divisions, we might expect them to be more interspersed with properly oriented periclinal divisions. If they are not misoriented periclinal divisions - could radial askew divisions be a distinctly oriented formative endodermal cell division? Two details hint at this, first, in addition to always being at the periphery of the endodermis, the daughters of radial askew endodermal divisions very often occur at the position where two cortex cells circumferentially meet (e.g. Figure 1C and 5A). Second, besides the endodermal cell division defects, the stele area of *irk* roots, particularly *irk-4*, is substantially wider than Wt due to increases in cell size (Campos et al., 2020; Goff et al., 2023). This likely increases mechanical force on the peripheral cell layers and cell division orientations are altered in response to changes in mechanical tension (Shapiro et al., 2015; Louveaux et al., 2016; Marhava et al., 2019). Thus, radial askew endodermal divisions could be a distinct and specific division orientation that generates precisely placed cells poised to invade the cortex layer in response to widening of internal root cell types.

We show extensive activity of *pCYCD6;1* in *irk* roots and, consistent with its reported activity in Wt, it is associated with formative GT divisions (Sozzani et al., 2010). However, *pCYCD6;1* activity also strongly associates with endodermal LADs, which are proliferative divisions. It is possible that excess *pCYCD6;1* activity in *irk* mutants simply reflects the high number of endodermal cell divisions occurring at any one time and a general defect in spatiotemporal control of cell cycle activity. Persistent expression of cell cycle genes after cytokinesis could explain the excess endodermal cell divisions observed in *irk* mutants. However, there is no increase in (proliferative) transverse anticlinal endodermal divisions that would lengthen the *irk* root meristem (Campos et al., 2020), nor are these divisions associated with *pCYCD6;1* activity. Because all of the excess divisions in *irk* are oriented such that they broaden the root’s radial axis and associated with *pCYCD6;1* activity, we propose that IRK represses a GT cell division program specific to the root’s radial axis with CYCD6;1 downstream of IRK in this pathway.

In developmental biology, cell divisions are parsed into two groups: formative and proliferative. Formative divisions are considered to require specialized cues to initiate and properly orient; in plants, this often occurs in the long axis of the cell and creates an additional cell layer with a distinct identity. In contrast, proliferative plant cell divisions tend to occur along the shortest path to create equal sized daughter cells of the same identity. It is straightforward to suggest that distinct mechanisms regulate proliferative and formative divisions. However, this is not very satisfying when considering the endodermal divisions linked to IRK and CYCD6;1. While neither IRK nor CYCD6;1 have known roles in proliferative cell divisions in the root’s longitudinal axis (Sozzani et al., 2010; Campos et al., 2020), they are linked to proliferative divisions in the radial axis. This may mean that distinctly oriented proliferative divisions are also differentially regulated from each other, in addition to the differential regulation between proliferative and formative divisions. However, a more parsimonious explanation may be that root cells separately regulate divisions that occur in different orientations. While this may seem more complex than the currently accepted formative vs. proliferative division paradigm, it may be more aligned with the underlying logic of plant body plan elaboration in respect to its organ axes. Regardless, our study of IRK indicates that a distinct pathway operates to repress cell divisions that specifically broaden the radial axis of the root GT - whether those divisions are proliferative or formative.

## MATERIALS & METHODS

### Plant material and growth conditions

Seeds were surface sterilized with chlorine gas and plated on MS agar media. The media contained 0.2X Murashige and Skoog salts (Caisson labs), 0.5g/L MES, 1% Sucrose, and 1% agar (Difco) and was at pH 5.7. Plates were sealed with parafilm and placed at 4°C for 24-72 hours for stratification. After stratification, plates were placed vertically in a Percival incubator and grown under 16 hrs light/8hrs dark at a constant temperature of 22°C. For each experiment, the genotypes being analyzed were grown side-by-side on the same plate. *Arabidopsis thaliana* ecotype Col-0 was used as the Wild type (Wt). The genotypes *irk-1, irk-4, irk-4 pIRK:IRKΔK:GFP*, and *irk-4 pIRK:IRK:GFP* were obtained as previously described (Campos et al., 2020; Rodriguez-Furlan et al., 2022). The cell type-specific reporters *pCO2:nlsYFP (Paquette and Benfey, 2005), pSCR:erGFP* (Di Laurenzio et al., 1996; Wysocka-Diller et al., 2000), and *pCYCD6;1::GUS:GFP* (Sozzani et al., 2010) were received from the Benfey lab and were crossed the *irk* mutants as previously described (Campos et al., 2020).

### Confocal microscopy and image analysis

Roots were imaged at day 6 post-stratification (dps) and stained with ∼10 μM propidium iodide (PI) solubilized in water for 1-2 minutes; then, imaged on a Leica SP8 upright microscope and imaging system housed in the Van Norman lab. Fluorescent protein (FP) signals were captured using the following settings: PI (excitation 536 nm, emission 585-660 nm) and GFP/YFP (excitation 488 nm, emission 492-530 nm). Z-stacks were acquired and analyzed with the orthogonal sectioning tool of the LASX software. The total number of endodermal divisions per root were counted from the QC to QC + 120μm or 20 cells as previously described (Campos and Van Norman, 2022). Representative images of roots expressing the *pCYCD6;1:GUS:GFP, pSCR:erGFP*, and *pCO2:nls:YFP* reporters were acquired from a transverse optical section at the 10^th^ endodermal cell (E10) shootward of the QC.

Signal separation GFP/YFP was performed by linear unmixing of a spectral image (Zimmermann et al., 2014), acquired with a inverted Zeiss 880 microscope with 32 channel spectral detector, objective 40X/1.2 W Korr FCS M27, MBS 488/561 and exciting with both 561 nm and 488 nm. Signal was collected simultaneously in 18 channels from 463 to 624 nm in ∼9 nm bands. GFP/YFP and PI signal was subsequently separated using Zeiss ZEN software linear unmixing tools.

For the images used to quantify endodermal cell divisions and generate the endodermal cell division map (see below), Z-stacks were acquired from each root at 512x512 pixels with 1 pm between optical sections. All three genotypes expressing a single reporter were grown on the same plate, 14-15 plants per genotype were imaged, and S10 plants per genotype were analyzed in detail. For each root, the optical sections were manually analyzed from the QC shootward 20 endodermal cells (E1-20), such that >160 endodermal cells per root were examined for excess endodermal cell divisions.

### Generation of the endodermal cell division maps

To generate the endodermal cell division maps for each genotype, we examined promoter activity and the presence of periclinal, radial askew, and longitudinal anticlinal divisions in each optical transverse section from just above the QC shootward to endodermal cell position 20 (E20). If a division was present, we documented whether the daughter cell was positive or negative for the reporter. For plants expressing *pCYCD6;1:erGFP*, we examined endodermal LADs, periclinal, and radial askew divisions. For plants expressing *pSCR:erGFP* and *pCO2:nls:YFP*, we only examined the endodermal periclinal and radial askew divisions, as the daughters of endodermal LADs remain endodermal cells. The total number of these divisions was quantified per root and then counted as GFP/YFP positive (+) or GFP/YFP negative (-). We note that the excess periclinal endodermal division phenotype in *irk-1* appears slightly repressed in the reporter backgrounds (compare Figure 1F to S1B, S2B, and S3B), however, this is likely due to general variability in the number of periclinal divisions in any genotype and relatively low number of roots (n = ∼40 vs. ∼10, respectively) examined per replicate.

The endodermal cell division map shows whether periclinal, radial askew, and/or longitudinal anticlinal divisions were present at a given position (E1-E20). This information is compressed as the map does not include the absolute number of divisions or map divisions in the root’s radial axis. Specifically, if a box in the division map is a solid color, then all the daughter cells for a given endodermal division that is present expressed the reporter - whether there was a single division or several. If some, but not all, of the daughter cells expressed the reporter, then the box in the map has a gradient fill. Finally, if none of the daughter cells expressed the reporter the box is unfilled (white). Endodermal division orientations are represented by different shapes in each box in the endodermal cell division maps: LADs are indicated by a black arrowhead on the right, internal side; periclinal divisions are indicated by a black arrowhead on the lower, internal side, and radial askew divisions are indicated by a curved line in the lower-right corner. If none of these endodermal cell divisions were detected, the box is empty.

### Assessing the spatial correlation between endodermal division orientations

To determine if endodermal radial askew divisions were correlated with periclinal or longitudinal anticlinal division, the endodermal cell division pattern along the root’s longitudinal axis was examined in each genotype expressing *pCYCD6;1:GUS:GFP*. In endodermal cell files from the QC upwards, transverse optical sections were examined for a radial askew division; then the adjacent shootward and rootward cells were examined for an abnormal division or no division. If the adjacent cell also had a radial askew division, we proceeded to look until an endodermal cell that had a periclinal or longitudinal anticlinal division or no division was found up to E20 for all cell files.

### Figure generation

Confocal images were exported from Leica software (LASX) as TIF files, which were cropped and resized in Adobe Photoshop. Graphs were generated with PRISM8 (GraphPad software https://www.graphpad.com/, San Diego, USA). Schematics were created in Adobe Illustrator. The figures containing confocal images, graphs, and schematics were assembled in Adobe Illustrator.

## Supporting information

All figures

## ACKNOWLEDGEMENTS

We thank former members of the Van Norman lab, including Dr. Jason Goff, Dr. Jessica N. Toth, and Dr. Cecilia Rodriguez Furlan, and Dr. Carolyn Rasmussen (UC, Riverside (UCR)) for discussions of and feedback on this manuscript while it was in preparation. We also thank Dr. Patricia Springer (UCR) for her insightful suggestion for how to display the endodermal division map. We appreciate access to the Zeiss 880 confocal microscope housed in the Institute of Integrative Genome Biology Microscopy Core Facility (UCR). This work was supported by funds from UCR, including Initial Complement (IC) and Academic Senate funds, USDA-NIFA-CA-R-BPS-5156-H, and NSF CAREER #1751385 awarded to J.M.V.N.

## AUTHOR CONTRIBUTIONS

Conceptualization: RC, JMVN, and RMIKR; Cell division methodology: RC; Endodermal division & FP mapping: RMIKR and RC; Dual FP examination: PPH; Resources: JMVN and RC.; Writing - Original Draft and Review and Editing: JMVN, RC, RMIKR, and PPH; Visualization: JMV, RMIKR, RC, and PPH; Supervision: JMVN; Funding Acquisition: JMVN.

## REFERENCES

van den Berg C, Willemsen V, Hage W, Weisbeek P, Scheres B (1995) Cell fate in the Arabidopsis root meristem determined by directional signalling. Nature 378: 62–65

Besson S, Dumais J (2011) Universal rule for the symmetric division of plant cells. Proc Natl Acad Sci U S A 108: 6294–6299

Campos R, Goff J, Rodriguez-Furlan C, Van Norman JM (2020) The Arabidopsis Receptor Kinase IRK Is Polarized and Represses Specific Cell Divisions in Roots. Dev Cell 52: 183–195.e4

Campos R, Van Norman JM (2022) Confocal Analysis of Arabidopsis Root Cell Divisions in 3D: A Focus on the Endodermis. Methods Mol Biol 2382: 181–207

Cui H (2016) Middle Cortex Formation in the Root: An Emerging Picture of Integrated Regulatory Mechanisms. Mol Plant 9: 771–773

De Smet I, Beeckman T (2011) Asymmetric cell division in land plants and algae: the driving force for differentiation. Nat Rev Mol Cell Biol 12: 177–188

Di Laurenzio L, Wysocka-Diller J, Malamy JE, Pysh L, Helariutta Y, Freshour G, Hahn MG, Feldmann KA, Benfey PN (1996) The SCARECROW gene regulates an asymmetric cell division that is essential for generating the radial organization of the Arabidopsis root. Cell 86: 423–433

Dolan L, Janmaat K, Willemsen V, Linstead P, Poethig S, Roberts K, Scheres B (1993) Cellular organisation of the Arabidopsis thaliana root. Development 119: 71–84

Facette MR, Rasmussen CG, Van Norman JM (2018) A plane choice: coordinating timing and orientation of cell division during plant development. Curr Opin Plant Biol 47: 47–55

Goff J, Imtiaz Karim Rony R, Ge Z, Hajný J, Rodriguez-Furlan C, Friml J, Van Norman JM (2023) PXC2, a polarized receptor kinase, functions to repress ground tissue cell divisions and restrict stele size. bioRxiv 2021.02.11.429611

Hartman KS, Muroyama A (2023) Polarizing to the challenge: New insights into polaritymediated division orientation in plant development. Curr Opin Plant Biol 74: 102383

Heidstra R, Welch D, Scheres B (2004) Mosaic analyses using marked activation and deletion clones dissect Arabidopsis SCARECROW action in asymmetric cell division. Genes Dev 18: 1964–1969

Livanos P, Müller S (2019) Division Plane Establishment and Cytokinesis. Annu Rev Plant Biol 70: 239–267

Louveaux M, Julien J-D, Mirabet V, Boudaoud A, Hamant O (2016) Cell division plane orientation based on tensile stress in Arabidopsis thaliana. Proc Natl Acad Sci U S A 113: E4294–303

Marhava P, Hoermayer L, Yoshida S, Marhavý P, Benková E, Friml J (2019) Re-activation of Stem Cell Pathways for Pattern Restoration in Plant Wound Healing. Cell 177: 957–969.e13

Muroyama A, Gong Y, Bergmann DC (2020) Opposing, Polarity-Driven Nuclear Migrations Underpin Asymmetric Divisions to Pattern Arabidopsis Stomata. Curr Biol. doi: 10.1016/j.cub.2020.08.100

Muroyama A, Gong Y, Hartman KS, Bergmann DC (2023) Cortical polarity ensures its own asymmetric inheritance in the stomatal lineage to pattern the leaf surface. Science 381: 54–59

Paquette AJ, Benfey PN (2005) Maturation of the ground tissue of the root is regulated by gibberellin and SCARECROW and requires SHORT-ROOT. Plant Physiol 138: 636–640

Pillitteri LJ, Guo X, Dong J (2016) Asymmetric cell division in plants: mechanisms of symmetry breaking and cell fate determination. Cell Mol Life Sci. doi: 10.1007/s00018-016-2290-2

Rasmussen CG, Bellinger M (2018) An overview of plant division-plane orientation. New Phytol 219: 505–512

Rasmussen CG, Humphries JA, Smith LG (2011) Determination of symmetric and asymmetric division planes in plant cells. Annu Rev Plant Biol 62: 387–409

Rasmussen CG, Wright AJ, Müller S (2013) The role of the cytoskeleton and associated proteins in determination of the plant cell division plane. Plant J 75: 258–269

Rodriguez-Furlan C, Campos R, Toth JN, Van Norman JM (2022) Distinct mechanisms orchestrate the contra-polarity of IRK and KOIN, two LRR-receptor-kinases controlling root cell division. Nat Commun 13: 235

Schaefer E, Belcram K, Uyttewaal M, Duroc Y, Goussot M, Legland D, Laruelle E, de Tauzia-Moreau M-L, Pastuglia M, Bouchez D (2017) The preprophase band of microtubules controls the robustness of division orientation in plants. Science 356: 186–189

Scheres B, Benfey PN (1999) ASYMMETRIC CELL DIVISION IN PLANTS. Annu Rev Plant Physiol Plant Mol Biol 50: 505–537

Shapiro BE, Tobin C, Mjolsness E, Meyerowitz EM (2015) Analysis of cell division patterns in the Arabidopsis shoot apical meristem. Proc Natl Acad Sci U S A 112: 4815–4820

Shiu SH, Bleecker AB (2001) Receptor-like kinases from Arabidopsis form a monophyletic gene family related to animal receptor kinases. Proc Natl Acad Sci U S A 98: 10763–10768

Shiu SH, Bleecker AB (2003) Expansion of the receptor-like kinase/Pelle gene family and receptor-like proteins in Arabidopsis. Plant Physiol 132: 530–543

Smertenko A, Assaad F, Baluška F, Bezanilla M, Buschmann H, Drakakaki G, Hauser M-T, Janson M, Mineyuki Y, Moore I, et al (2017) Plant Cytokinesis: Terminology for Structures and Processes. Trends Cell Biol 27: 885–894

Sozzani R, Cui H, Moreno-Risueno MA, Busch W, Van Norman JM, Vernoux T, Brady SM, Dewitte W, Murray JAH, Benfey PN (2010) Spatiotemporal regulation of cell-cycle genes by SHORTROOT links patterning and growth. Nature 466: 128–132

Van Norman JM (2016) Asymmetry and cell polarity in root development. Dev Biol 419: 165–174

Wallner E-S (2020) The value of asymmetry: How polarity proteins determine plant growth and morphology. J Exp Bot. doi: 10.1093/jxb/eraa329

Wysocka-Diller JW, Helariutta Y, Fukaki H, Malamy JE, Benfey PN (2000) Molecular analysis of SCARECROW function reveals a radial patterning mechanism common to root and shoot. Development 127: 595–603

Zimmermann T, Marrison J, Hogg K, O’Toole P (2014) Clearing Up the Signal: Spectral Imaging and Linear Unmixing in Fluorescence Microscopy. In SW Paddock, ed, Confocal Microscopy: Methods and Protocols. Springer New York, New York, NY, pp 129–148

