## Supplementary material for "Radial askew endodermal cell divisions reveal IRK functions in division orientation": All figures

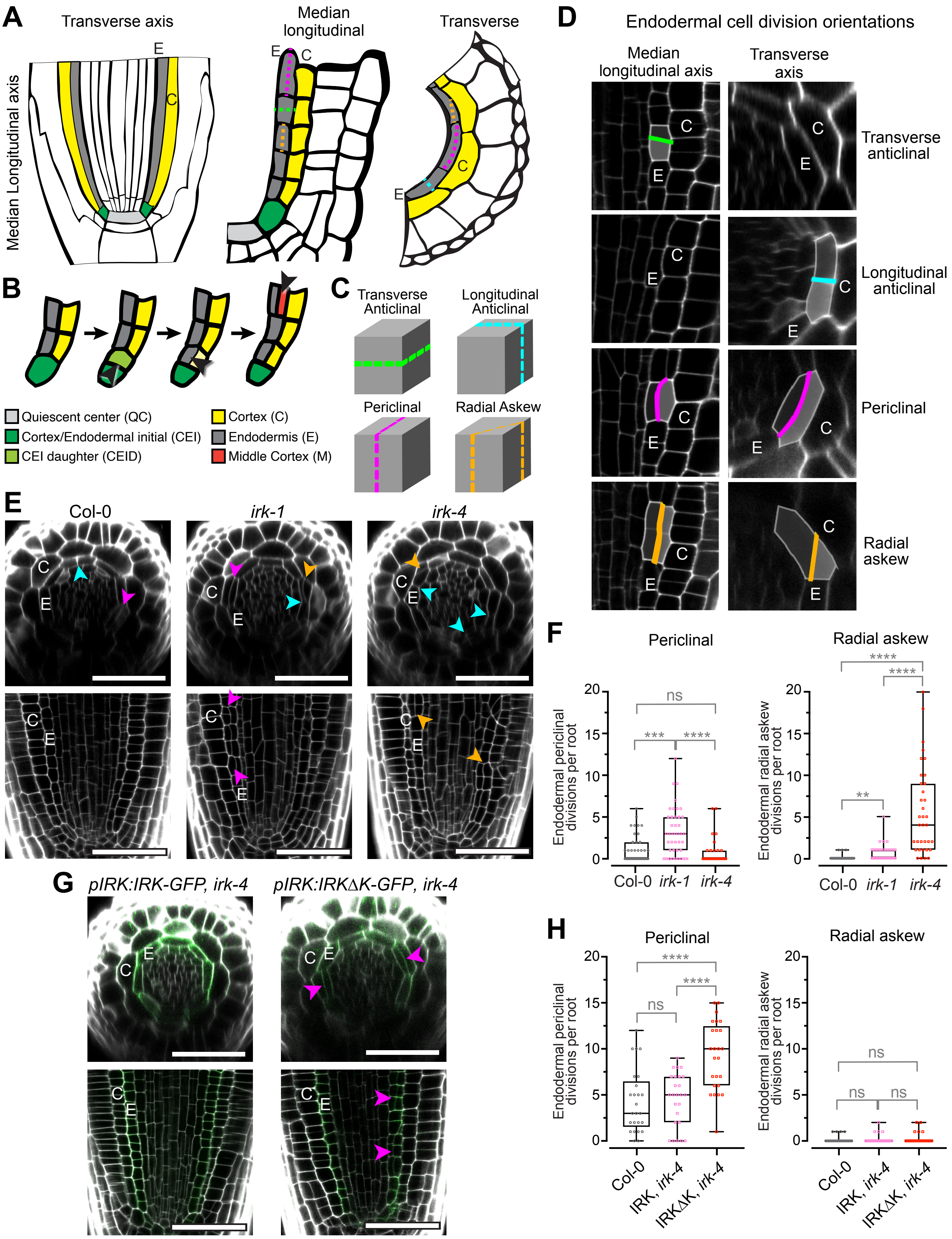

A
Transverse axis
Median Longitudinal axis
E
C
Median longitudinal
E
C
Transverse
C
E
B
Quiescent center (QC)
Cortex/Endodermal initial (CEI)
CEI daughter (CEID)
Cortex (C)
Endodermis (E)
Middle Cortex (M)
C
Transverse Anticlinal
Longitudinal Anticlinal
Periclinal
Radial Askew
D
Endodermal cell division orientations
Median longitudinal axis
Transverse axis
Transverse anticlinal
Longitudinal anticlinal
Periclinal
Radial askew
E
Col-0
irk-1
irk-4
F
Periclinal
Radial askew
Endodermal periclinal divisions per root
Endodermal radial askew divisions per root
Col-0
irk-1
irk-4
ns
***
****
**
G
pIRK:IRK-GFP, irk-4
pIRK:IRKΔK-GFP, irk-4
H
Periclinal
Radial askew
Col-0
IRK, irk-4
IRKΔK, irk-4

**Figure 1. Arabidopsis root cellular organization, cell division plane orientations, and *irk* phenotypes.** (A) Schematic of a root tip (left) with GT cell files and quiescent center (QC) highlighted. Close ups of (center) a median longitudinal and (right) transverse sections with endodermal division orientations indicated by dotted lines. (B) Schematic of ground tissue development from initial cell division to middle cortex formation. Black arrowheads indicate formative cell divisions. (C) Schematics of division orientations with single endodermal cells represented as cubes. (D) Confocal micrographs of a portion of the root tip showing the median longitudinal section (left) and transverse section (right) with endodermal cells that have undergone cell division. Note that transverse anticlinal divisions are not visible in the transverse axis and longitudinal anticlinal divisions are not visible in the longitudinal axis. (E) Confocal micrographs showing transverse sections (upper) and median longitudinal sections (lower) of root tips stained with propidium iodide (PI, gray) with excess endodermal cell divisions indicated by arrowheads: periclinal (magenta), longitudinal anticlinal (cyan), and radial askew (orange). (F) Quantification of the total number of periclinal and radial askew divisions per genotype (n = 35-40 roots per genotype). Graphs show the combined results from three replicates. (G) Confocal micrographs showing transverse sections (upper) and median longitudinal sections (lower) of *irk-4* expressing (left) *pIRK:IRK-GFP* and (right) *pIRK:IRK∆K-GFP* stained with PI (grey) merged with GFP (green). (H) Quantification of total number of periclinal and radial askew divisions in each genotype (n = 25 roots per genotype). Graphs show results combined from two replicates. Statistics: P-values indicated by asterisks: ** ($p \leq 0.01$), *** ($p \leq 0.001$), **** ($p \leq 0.0001$), and ns (not statistically significant) as determined by a one-way ANOVA using Dunn's multiple comparison test. Cell type abbreviations as indicated in panel (B) legend. Scale bars: 50 μm.

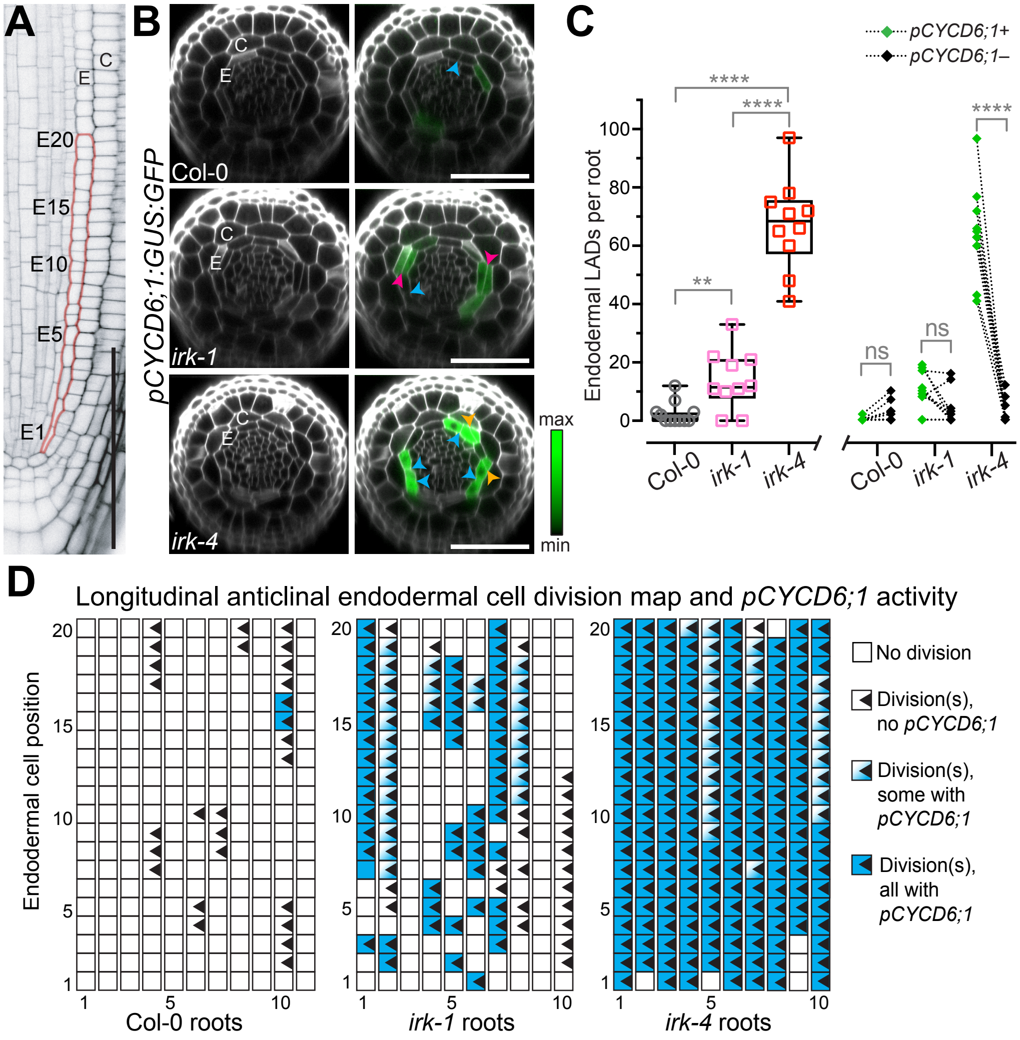


**Figure 2. Longitudinal anticlinal endodermal cell divisions in *irk* often exhibit *pCYCD6;1* activity**. (A) Confocal micrograph of a median longitudinal section of a portion of the root meristem stained with PI (gray scale) with red outline showing first 20 endodermal cells (E1-20) above the QC. (B) Confocal micrographs of transverse optical sections of Arabidopsis root meristems at E10. Roots were stained with PI (grey, left panels) to visualize cells and merged with *pCYCD6;1:GUS:GFP* (erGFP, green, right panels) activity. Endodermal cell divisions indicated by arrowheads: periclinal (magenta), longitudinal anticlinal (cyan), and radial askew (orange). Cell type abbreviations as indicated in Figure 1B. Scale bars: 50 µm. (C) Box plot (left) showing total number of endodermal LADs per root with whiskers indicating min/max with interquartile range and median shown with black boxes/lines, respectively, and colored symbols show measurements for individual roots. Paired point graph (right) showing LADs with (+) or without (–) *pCYCD6;1* activity. (D) Cell division map for endodermal LADs in the first 20 endodermal cells above the QC. Data shown are from one representative biological replicate of ≥ 2, 10-11 roots per genotype, with similar results. Statistics: ns = not statisticaly significant, p-values: ** < 0.01, **** < 0.0001, assayed by Mann-Whitney tests.

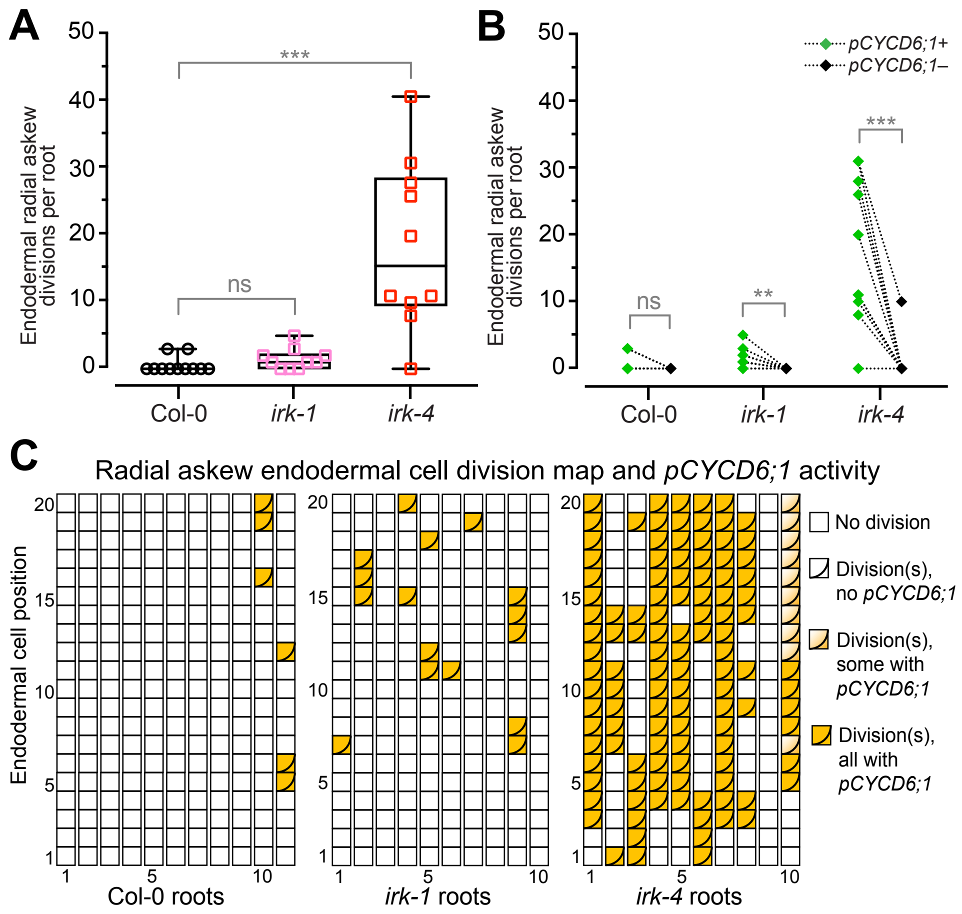


**Figure 3. Radial askew endodermal divisions consistently show *pCYCD6;1* activity**. (A, B) Quantification of radial askew endodermal divisions per root (the same roots examined as in Figure 2). (A) Box plot showing total number of radial askew divisions per root with whiskers indicating min/max with interquartile range and median shown with black boxes/lines, respectively, and colored symbols show measurements for individual roots. (B) Paired point graph showing the number of radial askew divisions with (+) or without (–) *pCYCD6;1* activity. (C) Cell division map for endodermal radial askew divisions in the first 20 endodermal cell above the QC (n = 10-11 roots per genotype). Data shown from one representative biological replicate of ≥ 2, with similar results. Statistics: ns = not statistically significant, p values: ** < 0.01, *** = 0.001, assayed by Mann-Whitney tests.

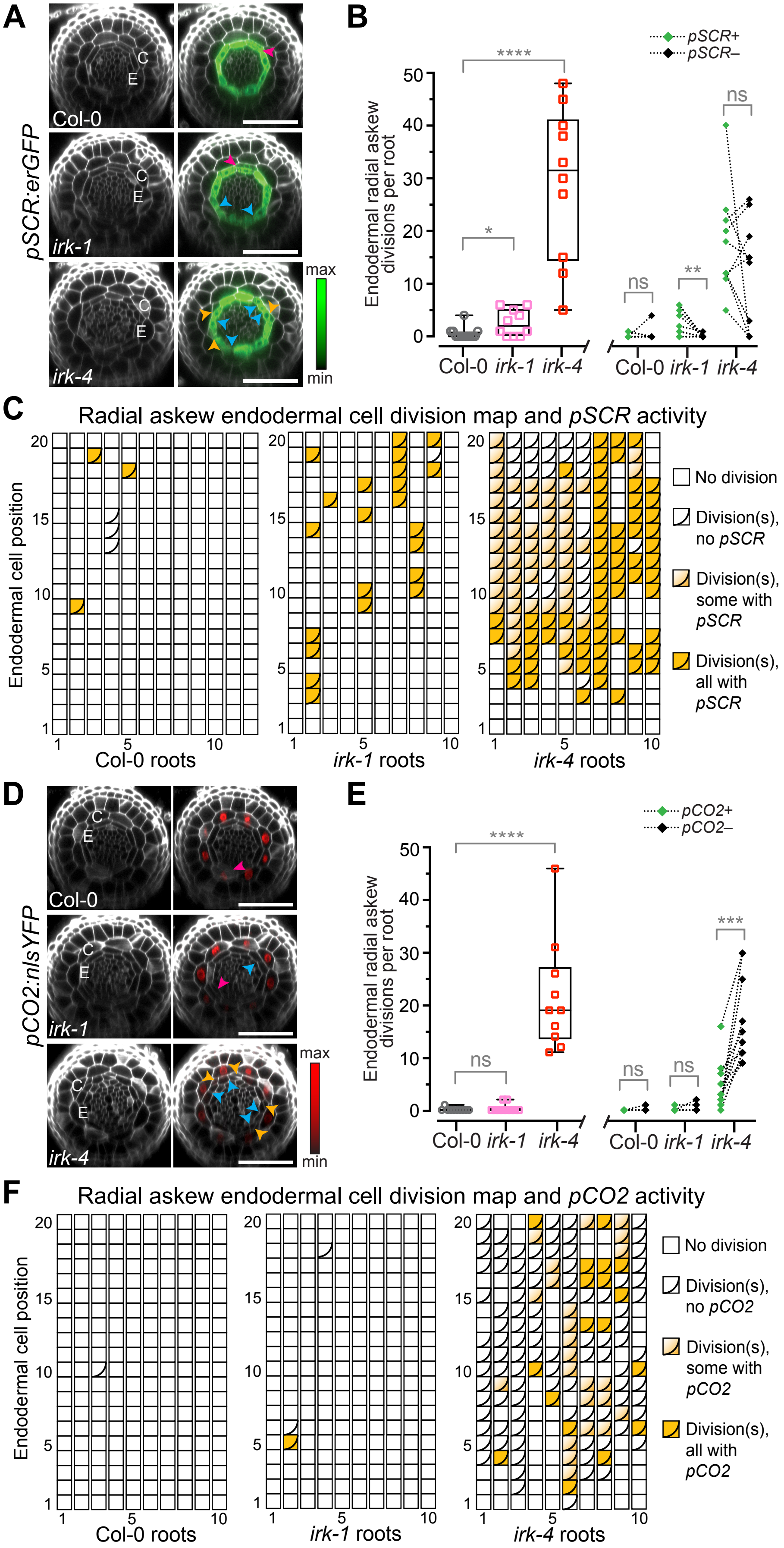
A
pSCR:erGFP
C
E
Col-0
irk-1
irk-4
max
min
B
pSCR+
pSCR–
Endodermal radial askew divisions per root
50
40
30
20
10
0
****
*
ns
**
ns
Col-0 irk-1 irk-4
Col-0 irk-1 irk-4
C
Radial askew endodermal cell division map and pSCR activity
Endodermal cell position
20
15
10
5
1
1
5
10
Col-0 roots
irk-1 roots
irk-4 roots
No division
Division(s), no pSCR
Division(s), some with pSCR
Division(s), all with pSCR
D
pCO2:nlsYFP
C
E
Col-0
irk-1
irk-4
max
min
E
pCO2+
pCO2–
Endodermal radial askew divisions per root
50
40
30
20
10
0
****
ns
ns
ns
***
Col-0 irk-1 irk-4
Col-0 irk-1 irk-4
F
Radial askew endodermal cell division map and pCO2 activity
Endodermal cell position
20
15
10
5
1
1
5
10
Col-0 roots
irk-1 roots
irk-4 roots
No division
Division(s), no pCO2
Division(s), some with pCO2
Division(s), all with pCO2

**Figure 4. Radial askew division daughters and expression of GT cell identity reporters**. (A, D) Confocal micrographs of transverse optical sections of Arabidopsis root meristems at position E10. Endodermal division orientations indicated by arrowheads: periclinal (magenta), longitudinal anticlinal (cyan), and radial askew (orange). Scale bars: 50 μm. (A) Images of roots expressing *pSCR:erGFP* stained with PI (grey, left) and merged with images of *pSCR* activity (erGFP, green, right). (B) Quantification of radial askew divisions per root and the numbers of radial askew divisions with (+) or without (-) *pSCR* activity. (C) Cell division map for endodermal radial askew divisions in the first 20 endodermal cells above the QC (n = 10-12 roots per genotype). (D) Images of roots expressing *pCO2:nlsYFP* were stained with PI (grey, left) and merged with images of *pCO2* activity (nlsYFP, red, right). (E) Quantification of radial askew divisions per root and the numbers of radial askew divisions with (+) or without (-) *pCO2* activity. (F) Cell division map for endodermal radial askew divisions (n = 10 roots per genotype). (B, E) Box plots show total number of radial askew endodermal divisions per root with whiskers indicating min/max with interquartile range and median shown with black boxes/lines, respectively, and colored symbols show measurements for individual roots. Paired point graphs showing the number of radial askew divisions with (+) *pSCR/pCO2* activity or without (–) *pSCR/pCO2* activity. Data shown are from one representative biological replicate of ≥2, with similar results. Statistics: ns = not statistically significant, p values: ** < 0.01, *** < 0.001, **** < 0.0001, assayed by Mann-Whitney tests.

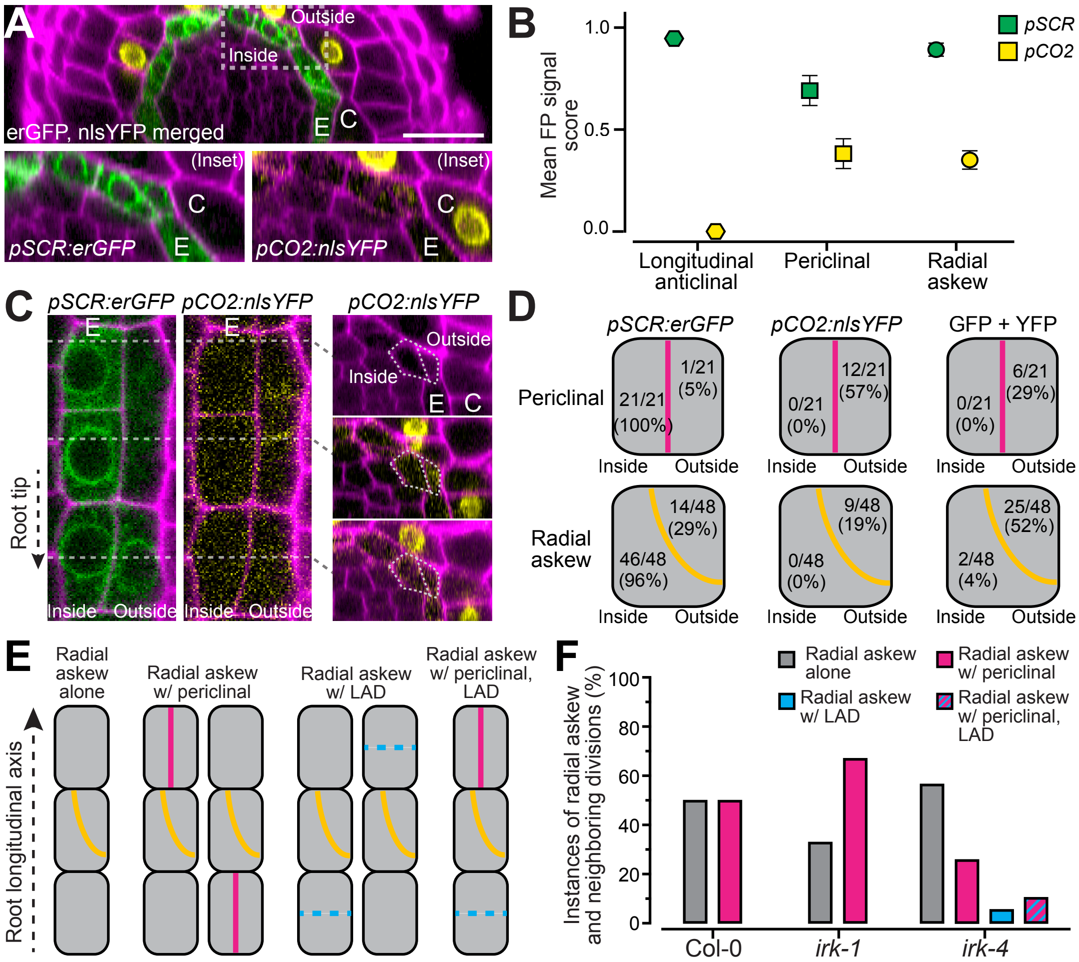


**Figure 5. The peripheral daughters of radial askew endodermal divisions acquire cortex cell identity.** (A) Confocal micrograph showing portion of a transverse section of an *irk-4* root meristem stained with PI (magenta) and simultaneously expressing reporters for endodermis (*pSCR:erGFP*, green) and cortex (*pCO2:nlsYFP*, yellow). Dashed rectangle indicates inset region (lower panels) showing a radial askew endodermal division with each reporter shown seperately. Scale bar: 25 µm. (B) Mean fluorescent signal score (based on subjective intensity) from cell identity reporters in the daughter cells of longitudinal anticlinal divisions (LADs, n = 44), periclinal divisions (n = 42), and radial askew divisions (n = 92). Obvious signal = 1, weak signal = 0.5, no signal = 0. (C) Longitudinal (left, center) and  transverse (right) sections showing progressive accumulation of *pCO2:nlsYFP* in the outer daughter cell of a radial askew division in the root's longitudinal axis. (D) Frequency of exclusive expression of cell identity markers (left, center) and incidence of their coexpression (right) in the daughter cells of periclinal and radial askew divisions (n = 9 roots). (E) Diagrams of endodermal cell files with a radial askew division (center cell, orange line) neighbored by cells that have not divided (left) or have undergone a periclinal (magenta line, left center) or longitudinal anticlinal division (LAD, cyan dashed line as these divisions are not visible in the longitudinal axis, right center) or both (right). (F) Bar graph showing instances of radial askew endodermal divisions with the neighboring cell states (as shown in E) as a proportion of the total instances per genotype (8 roots per genotype, Col-0: 2 instances; *irk-1*: 9 instances; *irk-4*: 16 instances). See methods section for additional details.

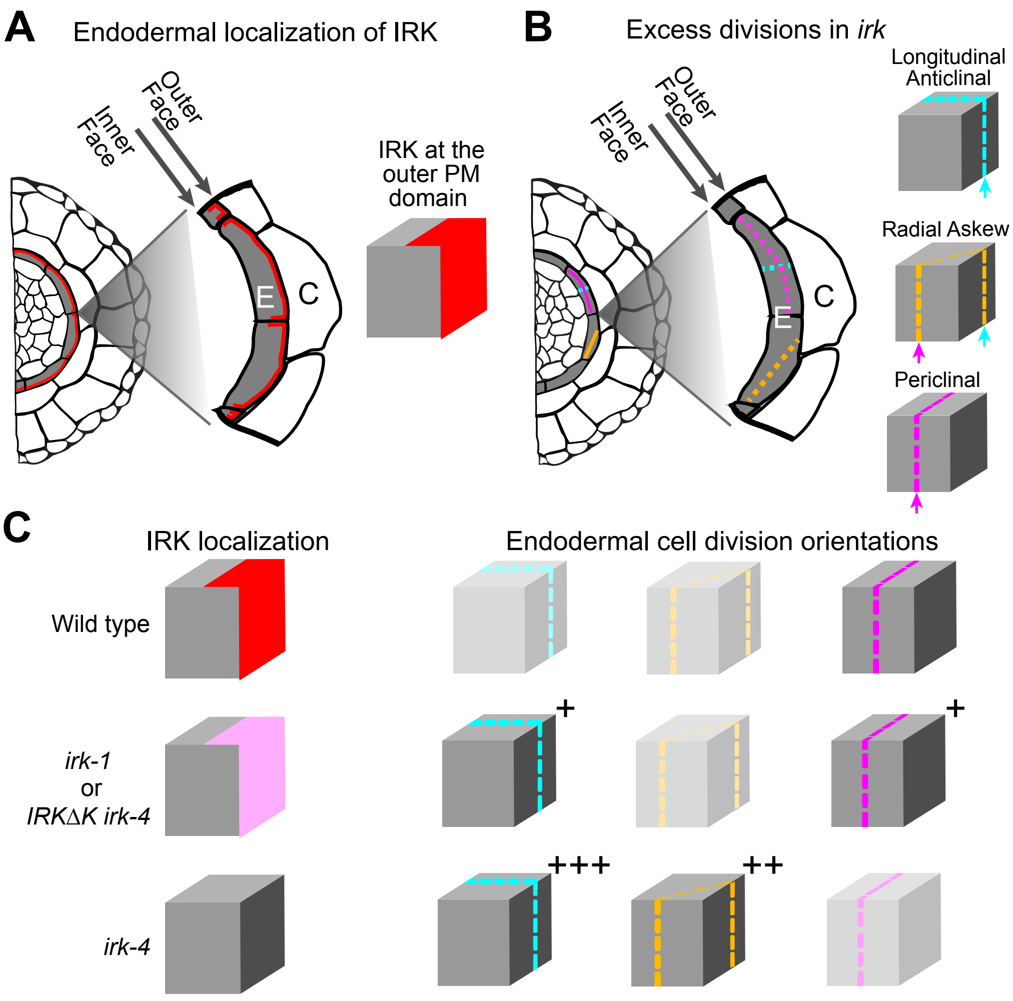


**Figure 6. Linking IRK polar localization to endodermal division orientations in *irk*.** Schematics of (A) IRK localization (red) at the outer lateral endodermal cell face and (B) excess endodermal division orientations in *irk*. (C) Schematics of single endodermal cells, represented as cubes, showing IRK localization (left) and endodermal division phenotypes (right) in the various genotypes. Plus (+) signs indicate the extent to which divisions in each of these orientations are present in *irk* compared to Wild type. The faded division orientations are rarely observed or, in the mutants, are not present at greater numbers than in wild type. Excess endodermal cell divisions in *irk-4* are frequently oriented such that the division plane contacts the endodermal cell face where IRK resides in Wild type, *irk-1*, and *irk-4* expressing IRKΔK-GFP.

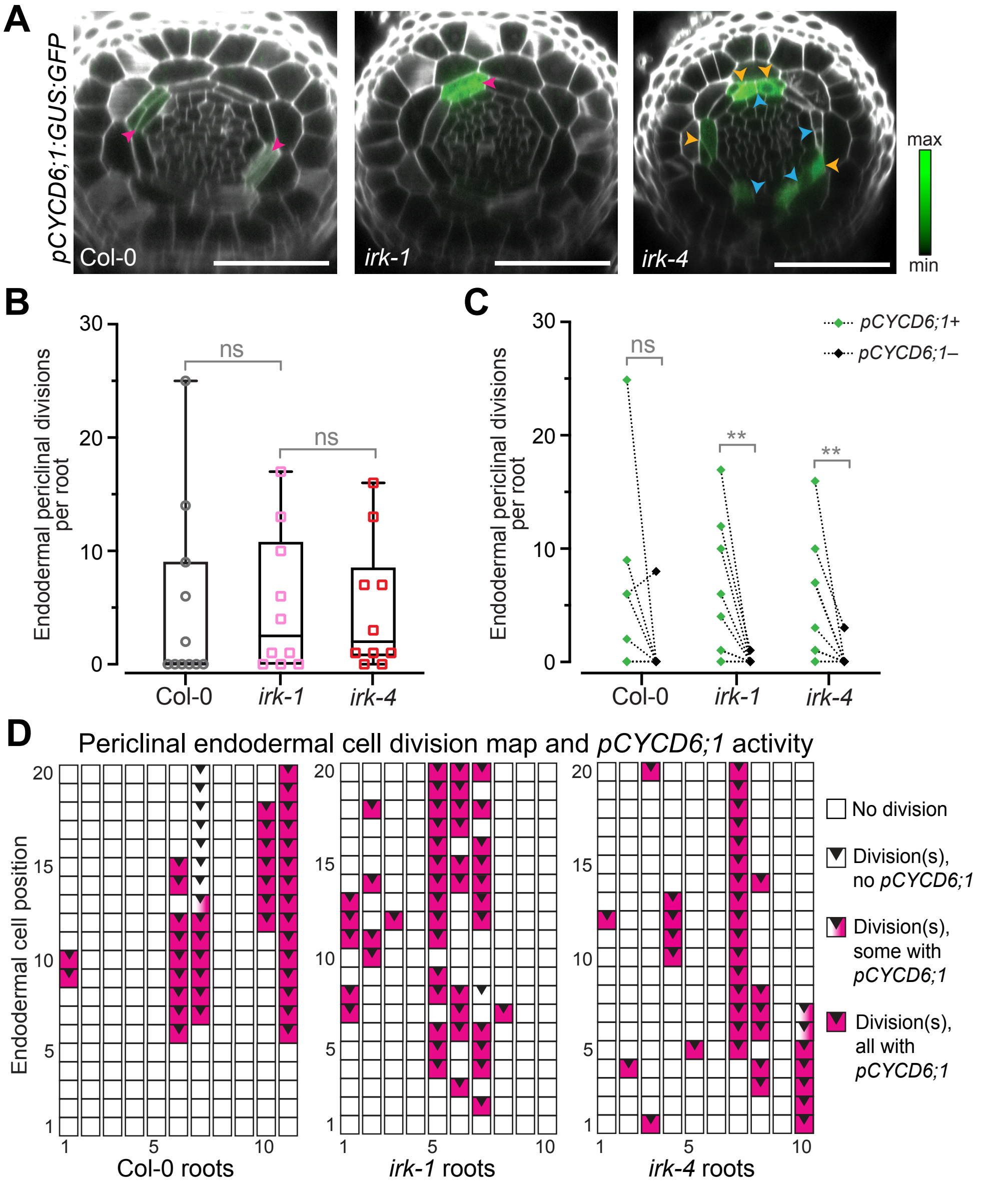


**Figure S1. Periclinal divisions are often associated with *pCYCD6;1* activity.** (A) Confocal micrographs of transverse optical sections of Arabidopsis root meristems expressing *pCYCD6;1:GUS:GFP*. Optical sections acquired at position E10 and the show PI channel (gray) merged with GFP (green). Scale bar = 50 µm. (B) Box plot showing total number of periclinal divisions per root with whiskers indicating min/max with interquartile range and median shown with black boxes/lines, respectively, with colored symbols showing measurements for individual roots. (C) Paired point graph showing the number of divisions with (+) or without (–) *pCYCD6;1* activity. (D) Cell division map for periclinal divisions in the first 20 endodermal cell above the QC (n = 10-11 roots per genotype, the same roots were analyzed in Figure 3). Data shown from a single representative biological replicate of ≥ 2, with similar results. Statistics: ns = not statistically significant, p values: ** < 0.01, assayed by Mann-Whitney test.

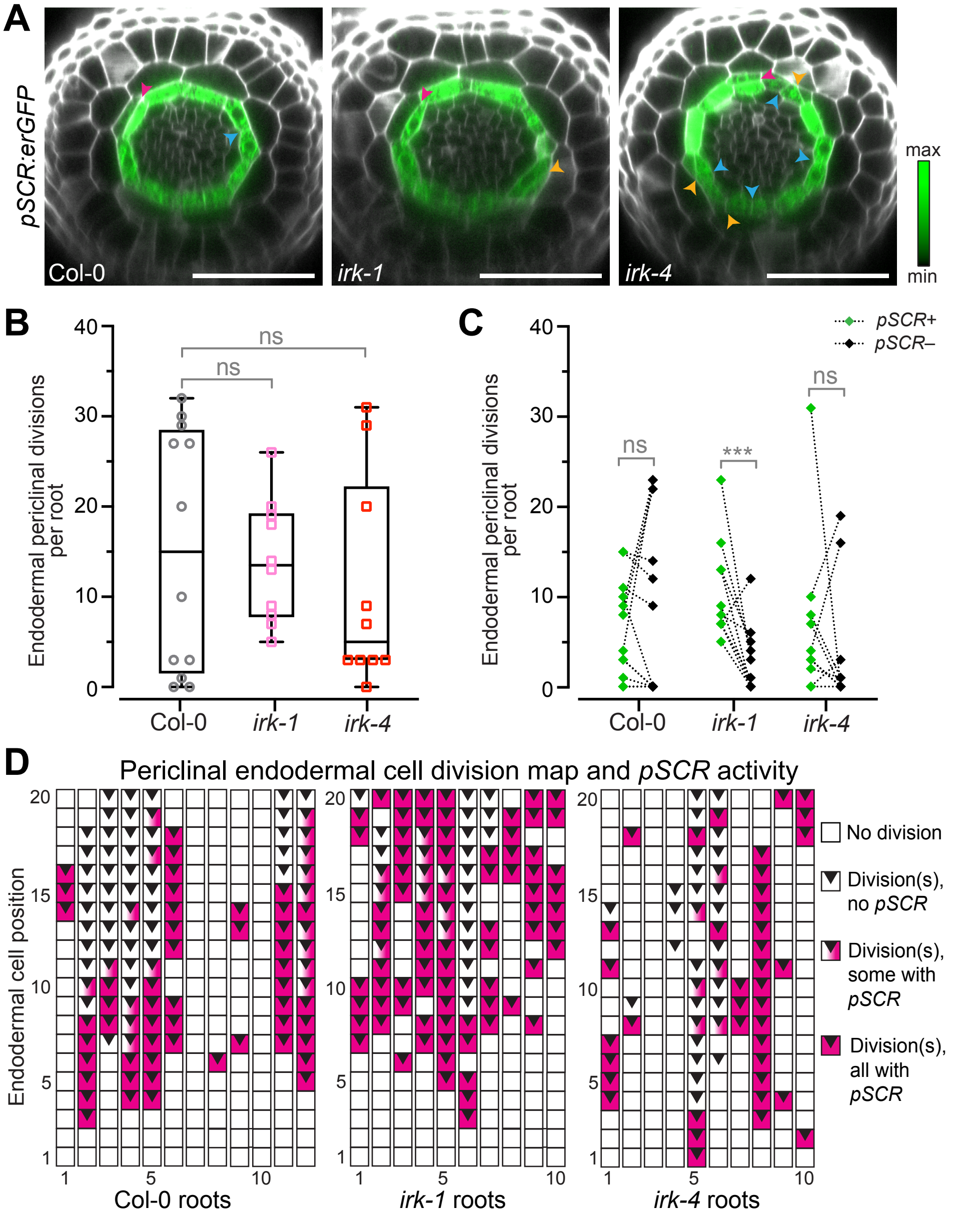


**Figure S2. Periclinal endodermal division daughter cells often show *pSCR* activity.** (A) Confocal micrographs of transverse optical sections of Arabidopsis root meristems expressing *pSCR:erGFP*. Optical sections acquired at position E10; images show PI channel (gray) merged with GFP (green). Scale bar = 50 µm. (B) Box plot showing total number of periclinal divisions per root with whiskers indicating min/max with interquartile range and median shown with black boxes/lines, respectively, and colored symbols show measurements for individual roots. (C) Paired point graph showing the number of divisions with (+) or without (–) *pSCR* activity. (D) Cell division map for periclinal endodermal divisions in the first 20 cells above the QC (n = 10 roots per genotype, the same roots were analyzed in Figure 4). Data shown are from a single representative biological replicate of ≥ 2, with similar results. Statistics: ns = not statistically significant, p values: *** < 0.001, as assayed by Mann-Whitney test.

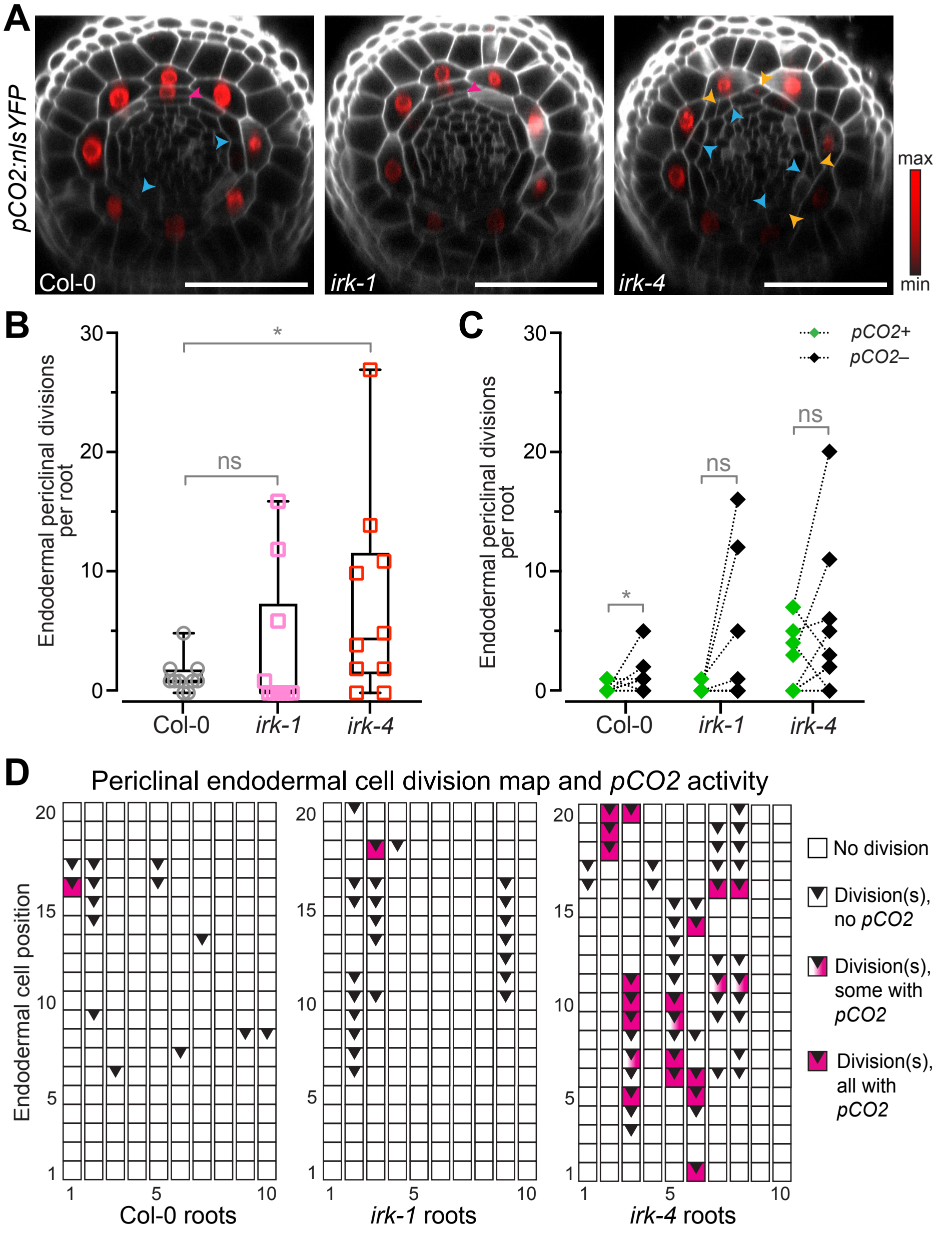


**Figure S3. Periclinal endodermal division daughter cells occasionally show *pCO2* activity.** (A) Confocal micrographs of transverse optical sections of Arabidopsis root meristems expressing *pCO2:nlsYFP*. Optical sections acquired at position E10 and images show PI channel (gray) merged with YFP (red). Scale bar = 50 µm. (B) Box plot showing total number of periclinal divisions per root with whiskers indicating min/max with interquartile range and median shown with black boxes/lines, respectively, and colored symbols show measurements for individual roots. (C) Paired point graph showing the number of divisions with (+) or without (–) *pCO2* activity. (D) Cell division map for periclinal divisions in the first 20 endodermal cell above the QC (n = 10 roots per genotype, the same roots were analyzed in Figure 4). Data shown are from a single representative biological replicate of ≥ 2, with similar results. Statistics: ns = not statistically significant, p values: * < 0.05, assayed by the Mann-Whitney test.
